# Loss of Resf1 reduces the efficiency of embryonic stem cell self-renewal and germline entry

**DOI:** 10.1101/2021.05.25.445589

**Authors:** Matúš Vojtek, Ian Chambers

## Abstract

Retroelement silencing factor 1 (RESF1) interacts with the key regulators of mouse embryonic stem cells (ESCs) OCT4 and NANOG, and its absence results in sterility of mice. However, the function of RESF1 in ESCs and germ line specification is poorly understood. In this study, we used *Resf1* knockout cell lines to determine the requirements of RESF1 for ESCs self-renewal and for *in vitro* specification of ESCs into primordial germ cell-like cells (PGCLCs). We found that deletion of *Resf1* in ESCs cultured in serum and LIF reduces self-renewal potential whereas episomal expression of RESF1 has a modest positive effect on ESC self-renewal. In addition, RESF1 is not required for the capacity of NANOG and its downstream target ESRRB to drive self-renewal in the absence of LIF. However, *Resf1* deletion reduces efficiency of PGCLC differentiation *in vitro*. These results identify Resf1 as a novel player in the regulation of pluripotent stem cells and germ cell specification.

## Introduction

Pluripotency is a feature of early embryonic epiblast and derivative cell lines (Martello and Smith, 2014). Pluripotent cells exist in naïve or primed states (Nichols and Smith, 2009), or an intermediate formative state (Kinoshita and Smith, 2018), from which cells directly differentiate into the germline. Of these pluripotency states, naïve embryonic stem cells (ESCs) are the best characterised (Martello and Smith, 2014). ESC identity is controlled by a gene regulatory network (GRN) centred around Oct4 (*Pou5f1*), Sox2 and Nanog. While Oct4 and Sox2 are uniformly expressed in all pluripotent states, Nanog expression is reduced during the formative state (Hayashi et al., 2011).

Both the germline and the naïve epiblast show dependencies on NANOG: constitutive *Nanog* deletion prevents specification of the naïve epiblast (Mitsui et al., 2003; Silva et al., 2009), while germline specific *Nanog* deletion reduces the number of PGCs in mid-gestation mouse embryos (Chambers et al., 2007; M. Zhang et al., 2018). On the other hand, Nanog overexpression sustains ESC self-renewal in the absence of the otherwise requisite leukemia inhibitory factor (LIF) (Chambers et al., 2003; Mitsui et al., 2003). Indeed, the level of NANOG expression determines the efficiency of ESC self-renewal, with *Nanog*^-/-^ ESCs having a reduced but residual self-renewal efficiency and *Nanog*^+/-^ ESCs having an intermediate self-renewal efficiency (Chambers et al., 2007). Nanog overexpression can also induce specification of germline competent epiblast-like cells (EpiLCs) into PGC-like cells (PGCLCs) *in vitro* without the otherwise requisite cytokines (Murakami et al., 2016). Similarities between the GRN of ESCs and PGCs are also highlighted by the capacity of the NANOG target gene, *Esrrb,* to maintain LIF-independent self-renewal in *Nanog^-/-^* ESCs and to restore wild type PGC numbers in mouse embryos where *Nanog* was specifically deleted from the germline (M. Zhang et al., 2018).

Nanog interacts with over 100 proteins in ESCs (Gagliardi et al., 2013). However, the requirements of these interactions for NANOG function and ESC self-renewal are largely unknown. Here we examine the function of the NANOG partner protein, RESF1 (also known as KIAA1551, GET) in ESC self-renewal and germ line specification. Our findings show that RESF1 has a modest positive effect on ESC self-renewal and that absence of RESF1 decreases efficiency of germ line specification.

## Results

### Resf1 deletion reduces ESC self-renewal and responsiveness of ESCs to LIF

Retroelement silencing factor 1 (RESF1) is a poorly characterised protein that interacts with the core pluripotency proteins OCT4 and NANOG in ESCs (Gagliardi et al., 2013; van den Berg et al., 2010). To study the function of RESF1 in mouse ESC self-renewal, we generated *Resf1^-/-^* ESCs using CRISPR/Cas9 (Figure 1A). We used two sets of four guide RNAs (gRNAs) positioned upstream of the transcription start site and downstream of the polyadenylation signal to delete the entire *Resf1* gene (Figure 1A). Wild type E14Tg2a ESCs were transfected with eSpCas9 plasmids encoding individual gRNAs, Cas9 and either green fluorescent protein (GFP) or mCherry, depending on whether the gRNA target site was 5’ or 3’ to the *Resf1* gene respectively (Figure 1A). Single cells expressing both GFP and mCherry were isolated and assessed for deletion of *Resf1* by PCR using two primer pairs (Figure 1A). Primer pair A amplifies a 680 bp sequence from the wild type *Resf1* intron IV (Figure 1A). If *Resf1* is deleted, primer pair A no longer amplifies a product. Primer pair B spans the entire *Resf1* locus (Figure 1A), and a large distance prevents PCR amplification from wild type cells under the reaction conditions used. However, when *Resf1* is deleted, these primers come into proximity to yield a product of ∼650bp. Two clones (c4 and c24) were identified in which the pattern of PCR amplification with primers A and B suggested deletion of the *Resf1* gene (Figure 1B). For c4, the presence of two bands of differing sizes suggested that each *Resf1* allele had been deleted using different gRNA pairs. Sequencing of the PCR products confirmed this and also revealed that each of the c24 *Resf1* alleles had undergone distinct deletion events (Figure S1).

**Figure 1:**
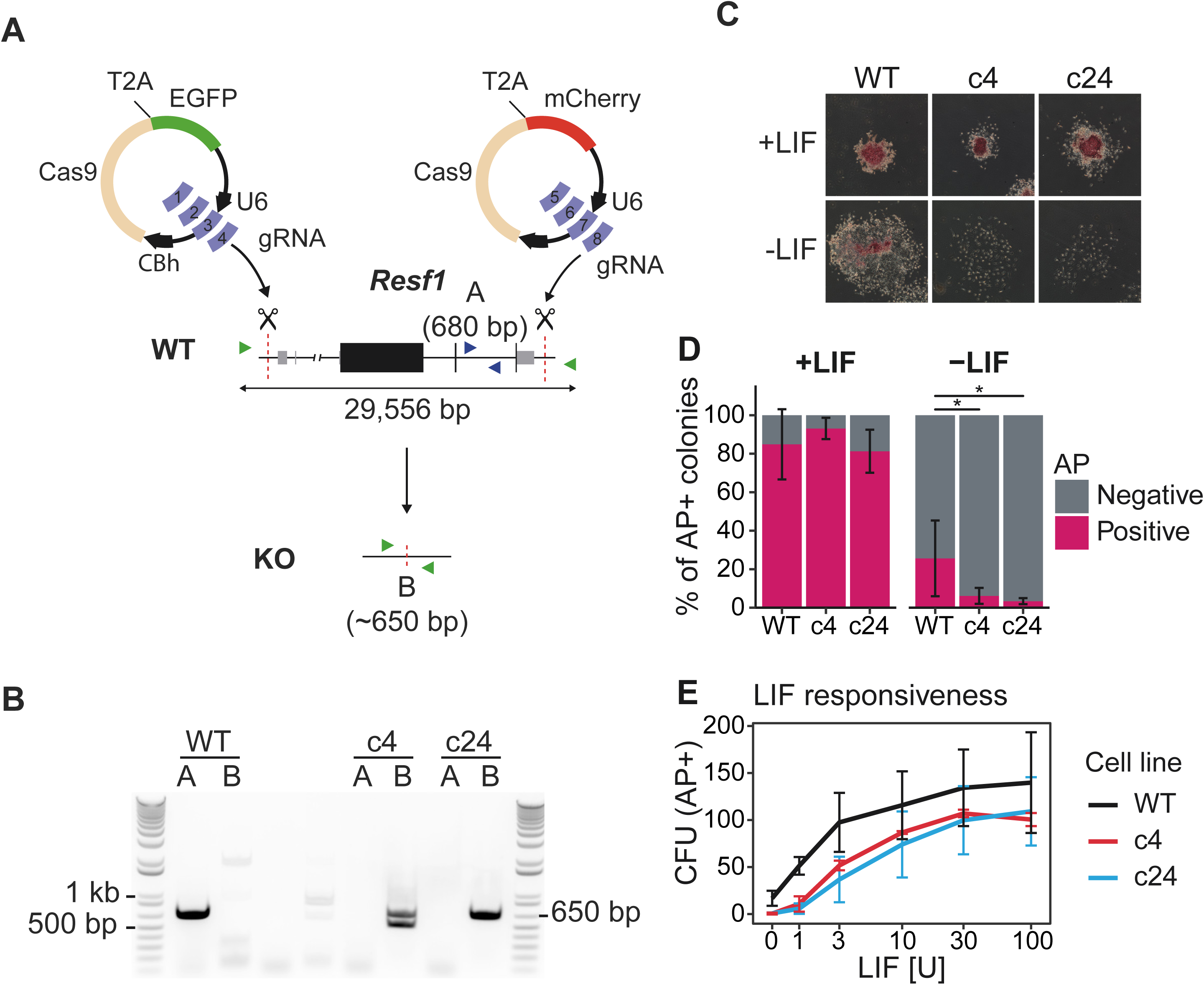
Deletion of *Resf1* decreases ESC self-renewal and responsiveness to LIF. **A)** Scheme of the deletion strategy used at the *Resf1* locus. The line diagram shows *Resf1*, with lines representing introns and non-transcribed regions, thick black boxes represent coding regions of exons and thin grey boxes represent UTRs. *Resf1* was deleted by targeting Cas9 via sets of gRNAs complementary to sites (red dotted lines) lying upstream of the transcription start site and downstream of the polyadenylation signal of the *Resf1* gene. Deletion of *Resf1* was assessed by PCR using primer pairs nested within intron IV (A, blue triangles) or flanking the targeted sites (B, green triangles). To delete *Resf1*, wild type (WT) ESCs were transfected with four CBh-eSpCas9-T2a-EGFP and four CBh-eSpCas9-T2a-mCherry plasmids carrying 8 distinct gRNAs. **B)** Electrophoresis of PCRs from WT ESCs and putative *Resf1^-/-^* ESC clones (c4 and c24) using primers A or B. **C-E,** Clonal ESC self-renewal assays. **C)** Representative images of the colonies formed by the indicated ESCs in the presence or absence of LIF. Colonies were stained for alkaline phosphatase (AP) 6 days after plating. **D)** The proportion of AP+ colonies formed by WT, *Resf1^-/-^* c4 and *Resf1^-/-^* c24 ESCs in the presence or absence of LIF. Bars represent mean ± SD (n = 5; * q < 0.05). **E)** Number of AP+ colony forming units (CFU) generated by WT or *Resf1*^-/-^ ESCs at different LIF concentrations (U/ml); mean ± SD (n = 4).

The capacity of *Resf1^-/-^* ESC clones 4 and 24 to self-renew in the presence or absence of LIF was examined after plating at clonal density. After 6 days in the presence of saturating levels of LIF, *Resf1^-/-^* ESCs formed colonies with a similar morphology to the parental wild type E14Tg2a ESCs (Figure 1C). In addition, the proportion of colonies expressing alkaline phosphatase (AP+) was also similar between the examined cell lines. In the absence of LIF, wild type cells do not produce any uniformly undifferentiated AP+ colonies but do yield a proportion of colonies containing differentiated and undifferentiated AP+ cells. The number of these mixed colonies was significantly reduced in both *Resf1^-/-^* clones (Wilcoxon rank-sum test, q < 0.05; Figure 1C, D). This suggests that *Resf1* deletion has a negative effect on ESC self-renewal.

To investigate this further, the colony-forming assay was repeated at decreasing concentrations of LIF. Both c4 and c24 *Resf1^-/-^* formed fewer AP+ colonies when cultured in LIF concentrations of 3 U/ml or less, with differences at higher LIF concentrations being less clear cut (Figure 1E, Figure S2A, B). This suggests that RESF1 sensitises the ESC response to low levels of LIF signalling.

### Deletion of *Resf1* decreases expression of pluripotency markers in ESCs cultured in serum

To investigate the basis of the reduced self-renewal efficiency of *Resf1^-/-^* ESCs, the expression of key pluripotency transcription factors was examined in *Resf1^-/-^* cells (Figure 2A). In both *Resf1^-/-^* ESC clones mRNA levels of *Nanog*, *Esrrb* and *Pou5f1* were reduced compared to the wild type ESCs, with *Esrrb* mRNA levels reduced by ∼4-fold (t-test, q < 0.05; Figure 2A). ESCs cultured in serum/LIF medium are heterogeneous for NANOG and ESRRB expression (Chambers et al., 2007; Festuccia et al., 2012). This heterogeneity can be eliminated in culture media containing two small inhibitors (2i) blocking FGF signalling and GSK3β (Silva and Smith, 2008; Ying et al., 2008). After switching to 2i/LIF culture, mRNA levels of *Esrrb, Nanog, Pouf51* or *Rex1* became equivalent in wild type and *Resf1^-/-^* ESCs (Figure 2B). This suggests that the reductions observed in serum/LIF may result from the cells initiating differentiation in serum/LIF culture.

**Figure 2:**
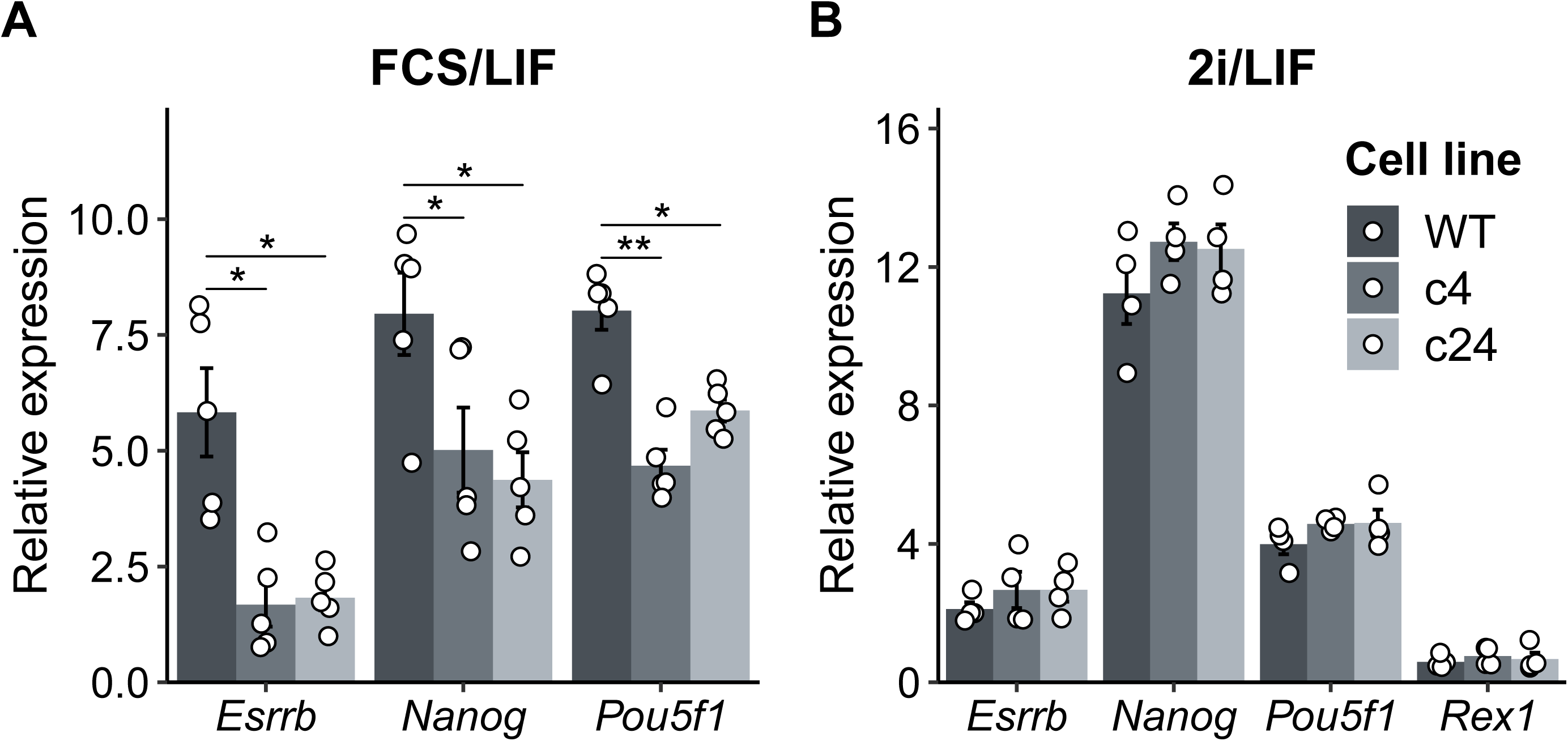
Effect of *Resf1* deletion on the expression of pluripotency markers in naïve pluripotent stem cells. mRNA levels of the indicated transcripts were determined in wild type (WT) and *Resf1*^-/-^ ESCs (c4 and c24) cultured in (**A**) serum/LIF medium (n = 5), (**B)** 2i/LIF medium (n = 4). Bars and whiskers represent mean ± SEM (n = 4). Scatter plots represent individual data points. * q < 0.05.

### Episomal expression of RESF1 has a modest positive effect on ESC self-renewal

As *Resf1* deletion reduced ESC self-renewal efficiency and decreased expression of pluripotency transcription factors, we hypothesised that enforced RESF1 expression may increase ESC self-renewal. To test this, Flag-RESF1 was expressed from an episome in *Resf1^+/+^* E14/T ESCs (Chambers et al., 2003) (Figure 3A). Transfected cells were cultured in selection medium at clonal density in the presence or absence of LIF for 8 days (Figure 3A), stained for alkaline phosphatase activity and quantified (Figure 3B, C). In the absence of LIF, NANOG conferred LIF-independent ESC self-renewal (Chambers et al., 2003), but RESF1 did not (Figure 3C). In contrast, in the presence of LIF, ESCs expressing episomal Flag-RESF1 formed significantly more AP+ colonies than ESCs transfected with an empty vector (Wilcoxson rank-sum test, q < 0.01), though fewer than obtained following Nanog transfection (Figure 3 C).

**Figure 3:**
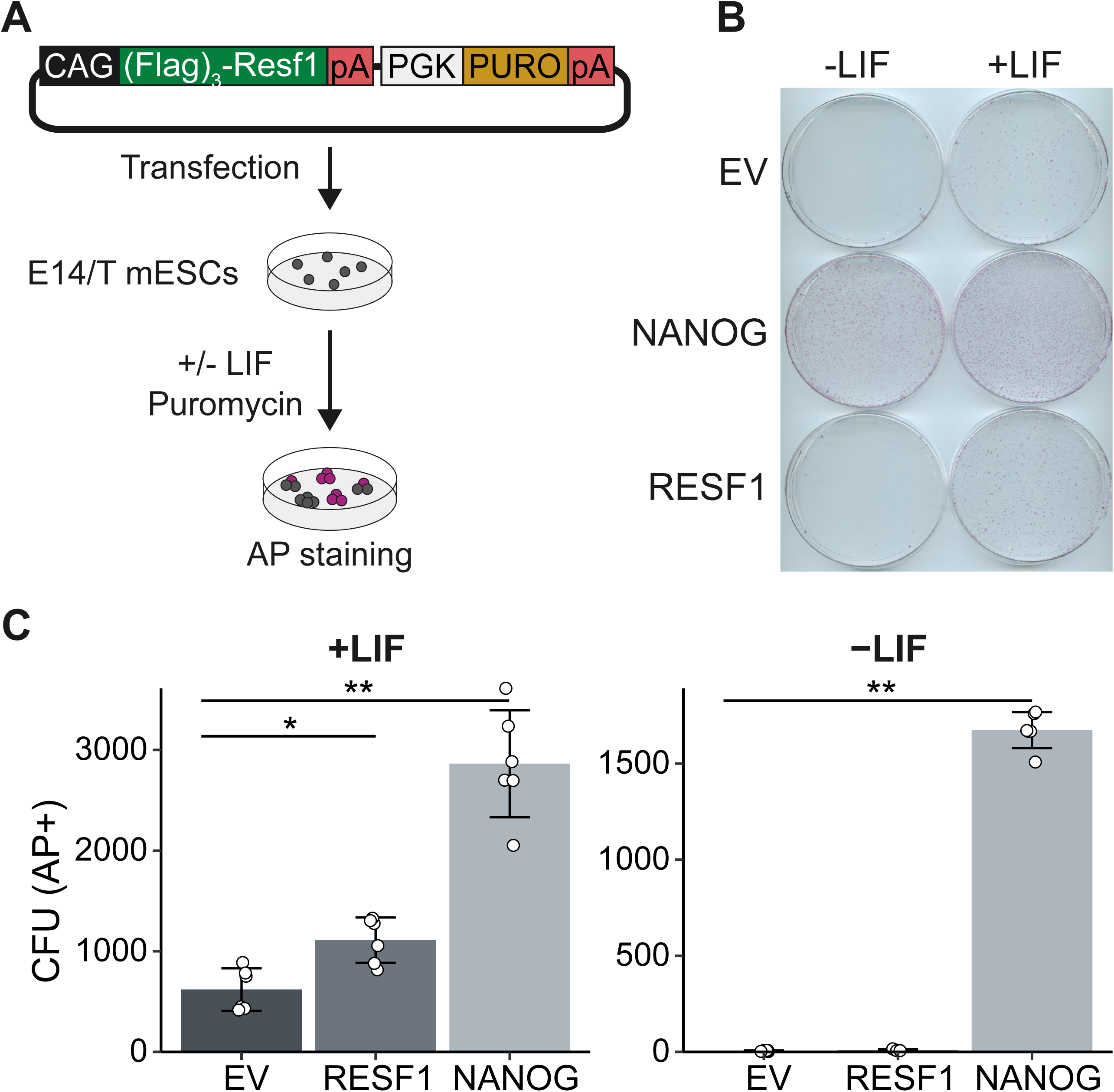
Effect of episomal expression of *Resf1* on ESC self-renewal. **A)** Strategy to assess the effect of episomal expression of RESF1 on ESC self-renewal. E14/T ESCs were transfected with the plasmid shown and cultured in the presence of puromycin in medium with or without LIF**. B)** After 8 days, colonies were stained for alkaline phosphatase (AP). Empty vector (EV) and a plasmid encoding Nanog in place of Resf1 provided controls. **C)** Quantification of colony numbers from **B.** Bars represent mean ± SD (n= 6) and scatter plots individual data points; ** q < 0.01, * q < 0.05.

### *Resf1* is not required for NANOG or ESRRB function in ESC self-renewal

The capacity of ESCs to self-renew is sensitive to NANOG levels (Chambers et al., 2007, 2003; Mitsui et al., 2003). As NANOG interacts with RESF1 (Gagliardi et al., 2013), we investigated the importance of RESF1 for NANOG function in ESC self-renewal. First, we validated the interaction between NANOG and RESF1 by co-immunoprecipitation. Episomally expressed Flag-RESF1 was immunoprecipitated from nuclear extracts of ESCs transfected with plasmids encoding Flag-RESF1 and HA-NANOG. When both Flag-RESF1 and HA-NANOG were co-expressed, Flag immunoprecipitation co-purified HA-NANOG (Figure 4A). In contrast, Flag antibody did not purify HA-NANOG from the control sample lacking Flag-RESF1 (Figure 4A). This confirms that RESF1 and NANOG interact in ESCs.

**Figure 4:**
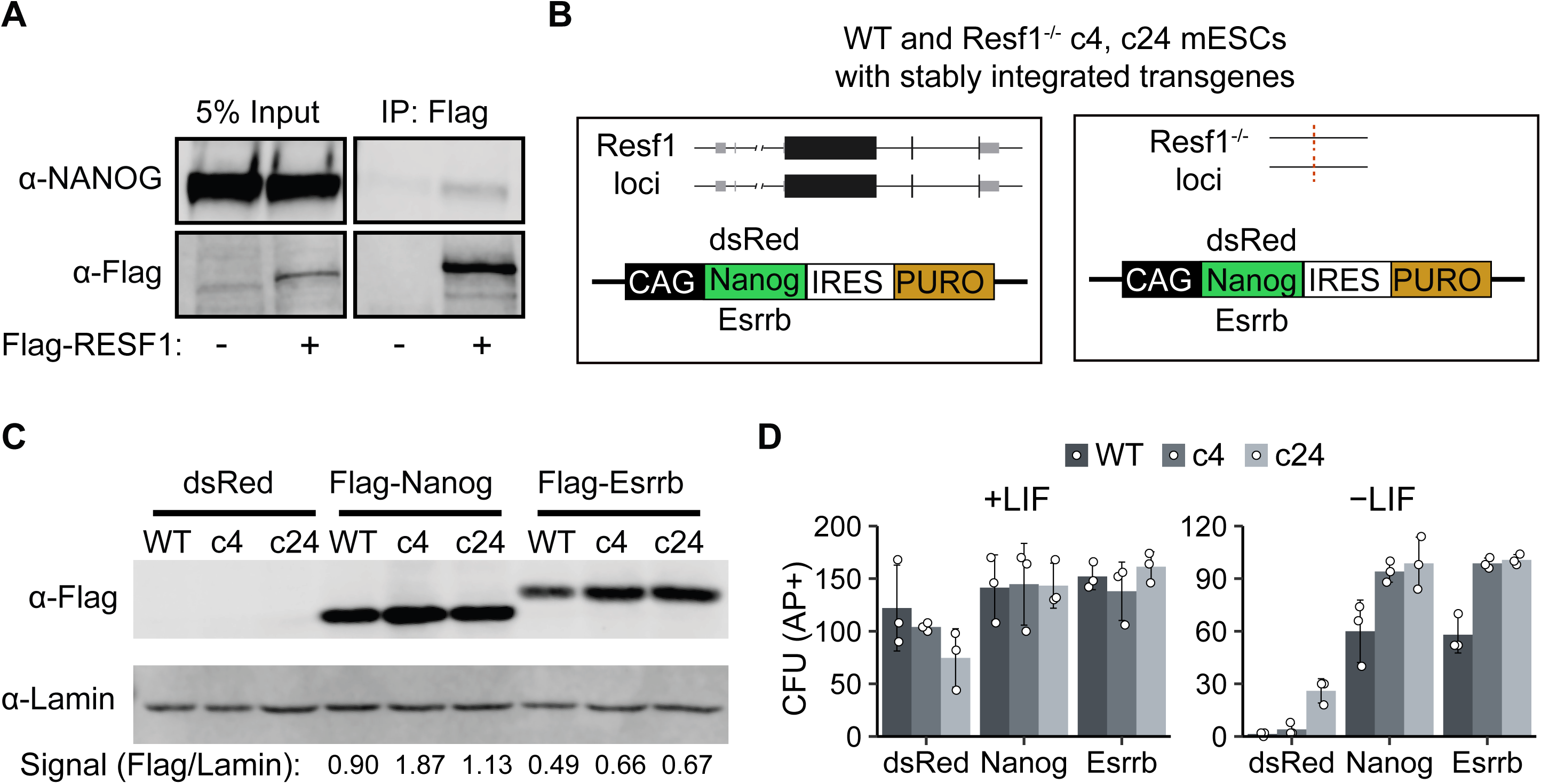
*Resf1* is dispensable for NANOG and ESRRB function to sustain LIF-independent self-renewal. **A)** Co-immunoprecipitation of Flag-RESF1 and NANOG from nuclear extracts of ESCs episomally expressing NANOG alone (-) or NANOG plus Flag-RESF1 (+). **B)** Schematic representation of wild type (WT) or *Resf1^-/-^* ESCs with stably integrated transgenes in which the puromycin resistance gene (PURO) is linked in the same transcript to either dsRed, Flag-Nanog or Flag-Esrrb. Black boxes represent coding exons, grey boxes represent non-coding exons. **C)** Immunoblot analysis of Flag expression after stable integration of dsRed, Flag-Nanog or Flag-Esrrb expression cassettes in WT and *Resf1^-/-^* ESCs (c4, c24); anti-LAMIN was used as a loading control. Relative Flag signal over LAMIN control is shown below. **D)** Quantification of alkaline positive (AP+) colony forming units (CFUs) formed by cells described in **B)** after 8-day culture in the presence or absence of LIF. Bars represent mean ± SD (n=3).

To examine the importance of RESF1 for NANOG function, we assessed whether *Resf1* was required for NANOG, or its downstream target ESRRB to confer LIF-independent self-renewal. Constitutive transgenes expressing Flag-NANOG-IRES-Puro, Flag-ESRRB-IRES-Puro or dsRed-Puro transgenes were stably integrated into wild type and *Resf1^-/-^* ESCs (Figure 4B). After 12-day selection, cell populations transfected with Nanog or Esrrb transgenes expressed Flag-NANOG or Flag-ESRBB (Figure 4C). The self-renewal capacity of these cell lines was next assessed by quantification of colony forming assays. In the presence of LIF, both wild type and *Resf1^-/-^* cells formed similar numbers of alkaline phosphatase positive colonies (Figure 4D, S2C). In the absence of LIF, expression of Flag-NANOG or Flag-ESRRB in wild type ESCs supported clonal ESC self-renewal (Figure 4D). Although overexpression of Flag-NANOG or Flag-ESRRB in *Resf1*^-/-^ cells appeared to increase alkaline phosphatase positive colony formation relative to wild type ESCs (Figure 4D), this could be due to a higher expression of NANOG and ESRRB transgenes in *Resf1^-/-^* cells (Figure 4C). These results indicate that *Resf1* is not required for ESRRB and NANOG to sustain LIF-independent self-renewal.

### Epitope tagging of endogenous *Resf1*

As there are no available antibodies for mouse RESF1, we generated ESC lines carrying an epitope tagged endogenous *Resf1* gene to facilitate the study of its molecular properties. To do this, we transfected E14Tg2a ESCs with a modification construct and a plasmid carrying a single gRNA and Cas9. The modification construct extended the *Resf1* open reading frame to include three V5 epitope tags, followed by an internal ribosomal entry site-green fluorescent protein cassette (IRES-GFP) and a single loxP site (Figure 5A).

**Figure 5:**
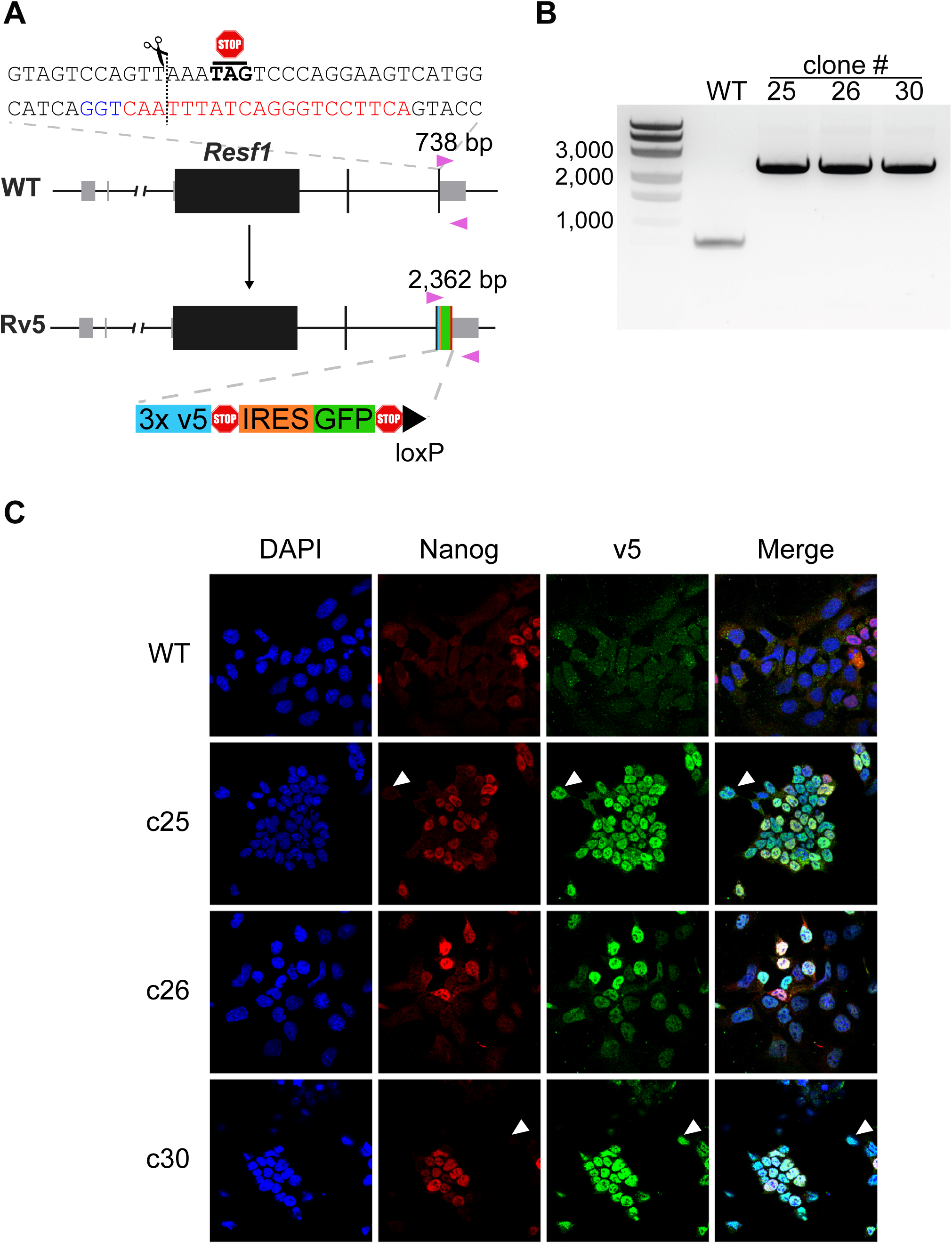
Epitope tagging of the endogenous *Resf1* gene. **A)** Cartoon representation of *Resf1* in wild type (WT) and Resf1-tagged (Rv5) cells. Lines represent introns and non-transcribed regions; thick boxes represent coding exons; thin grey boxes represent non-codding exons. Resf1 was tagged by inserting 3x v5-tag epitopes, stop codon, internal ribosome entry site (IRES), green fluorescent protein (GFP), stop codon and a loxP site in front of the Resf1 stop codon using CRISPR/Cas9. The complementary sequence of the gRNA (red) with the PAM sequence (blue) is shown. The expected cleavage site is indicated by a dotted line close to the *Resf1* stop codon. Genotyping primers (pink triangles) and expected PCR product sizes are shown. **B)** Genotyping of WT and Rv5 clones using primers flanking the insertion site. **C)** Immunostaining of WT and Rv5 clones for RESF1-v5 and Nanog. White arrows indicate cells expressing v5 but no NANOG.

After two days, single cells expressing high levels of GFP were isolated, expanded and genotyped. We used a primer pair flanking the insertion site that produces a 738bp band from the wild type allele and that increases in size to 2362bp upon insertion of the tagged modification (Figure 5A). Three ESC lines with an extended *Resf1* allele and lacking a wild type allele were identified (Figure 5B). Immunofluorescence confirmed expression of the v5 epitope in all three cell lines (Figure 5C). RESF1-v5 localized to the nucleus (Figure 5C) consistent with the previous observations using overexpression of RESF1 (Fukuda et al., 2018). Like NANOG, RESF1-v5 is expressed heterogeneously in ESCs cultured in serum/LIF. However, RESF1 marks a larger subset of ESCs than NANOG, with several NANOG negative cells express RESF1-v5 (Figure 5C, white arrows). This is similar to findings for NANOG-TET2 co-expression pattern (Pantier et al., 2019).

### Deletion of *Resf1* decreases efficiency of PGCLC specification

NANOG is required to provide wild type numbers of primordial germ cells (PGCs) *in vivo* (M. Zhang et al., 2018). In addition, enforced expression of NANOG in epiblast-like cells (EpiLCs) enables cytokine independent differentiation of PGC-like cells (PGCLCs) *in vitro* (Murakami et al., 2016; M. Zhang et al., 2018). RESF1 is reported to be required for fertility in mice, although the mechanism responsible is unknown (Dickinson et al., 2016). Therefore, we investigated the function of RESF1 in early germline specification.

First, we examined *Resf1* mRNA levels during germline specification *in vitro*. We quantified *Resf1* mRNA levels in wild type E14Tg2a naïve ESC cultures (serum/LIF and 2i/LIF medium), formative EpiLCs, primed epiblast stem cells (EpiSCs) and PGCLCs. *Resf1* mRNA levels in ESCs, EpiLCs and EpiSCs were similar, although median *Resf1* expression was higher when ESCs were cultured in serum/LIF medium (Figure S3A). Interestingly, day 4 PGCLCs expressed higher levels of *Resf1* than EpiSCs.

To investigate *Resf1* expression further, we analysed published single cell RNA sequencing datasets from mouse epiblast and PGCs between embryonic days (E) 6.5 and 8.5 (Pijuan-Sala et al., 2019). *Resf1* expression was detected in epiblast cells between E6.5 and E7.75 (Figure S3B). *Resf1* expression in epiblast cells is highest at E6.75 and decreases at later stages (Figure S3C). *Resf1* is also continuously expressed in PGCs between E6.75 to E8.5 (Figure S3B). In agreement with our Q-RT-PCR results, *Resf1* expression is higher in PGCs than in cells of post-implantation epiblast (Figure S3C). These results suggest that RESF1 might function in early germ cell development.

Therefore, we examined the capacity of *Resf1^-/-^* ESCs to specify PGCLCs *in vitro* (Figure 6A). Initially, wild type and *Resf1*^-/-^ cells formed EpiLCs expressing similar levels of EpiLC markers *Fgf5, Otx2* and *Pou3f1* (Figure S4A), indicating proper transition of *Resf1^-/-^* ESCs into PGC competent EpiLCs. Further aggregation of the wild type EpiLCs in the presence of PGC-specifying cytokine cocktail for 4 days induced surface expression of CD61 and SSEA1, which jointly mark PGCLCs (Hayashi et al., 2011) (Figure 6B, 6C, S5B). However, the proportion of SSEA1+CD61+ cells in the population was reduced in both clonal R*esf1^-/-^* cell lines (Figure 6B, C). Moreover, expression of mRNAs encoding the key PGC transcription factors *Ap2*γ*, Blimp1* and *Prdm14* was reduced in *Resf1^-/-^* cells (Figure 6D). This effect was clearest for *Blimp*1, which mRNA levels were lowered to ∼40% of wild type expression in both *Resf1^-/-^* clonal cell lines (t-test, q < 0.05; Figure 6D). Together these results indicate that *Resf1* contributes to efficient PGCLC differentiation *in vitro* and that the *Resf1* requirement may occur downstream of Ap2γ and upstream of *Blimp1* and CD61/SSEA1 expression.

**Figure 6:**
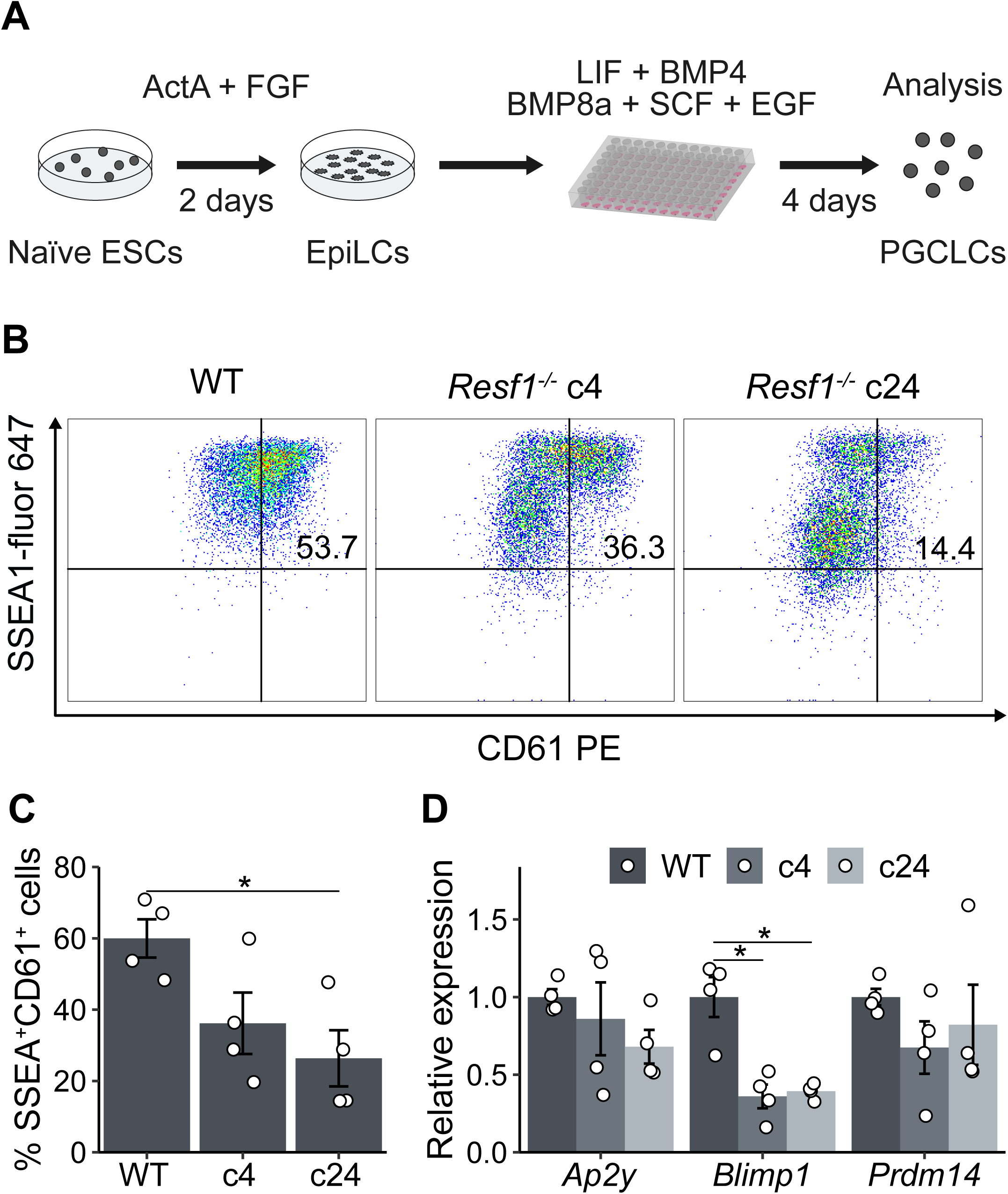
Effect of *Resf1* deletion on PGCLC differentiation. **A)** Scheme of differentiation of naïve ESCs into PGCLCs. ESCs are treated with Activin A and FGF2 for 2 days to form EpiLCs. EpiLCs are then aggregated in the presence of the indicated cytokines. **B)** Representative scatter plots of SSEA1 and CD61 expression measured by flow cytometry after 4 days of PGCLC differentiation using the indicated cell lines. Numbers represent percentage of CD61+SSEA1+ population. **C)** Quantification of CD61+SSEA1+ cell populations shown in **B** (n= 4). **D)** Relative expression of the indicated PGC markers in WT or *Resf1^-/-^* cells after 4 days of PGCLC differentiation (n=4). Bars represent mean ± SEM. Scatter plots represent individual data points. * q < 0.05.

## Discussion

RESF1 is a poorly characterised protein lacking known functional domains. However, the interaction of RESF1 with the pluripotency transcription factors NANOG and OCT4 (Gagliardi et al., 2013; van den Berg et al., 2010) suggests that RESF1 may function in ESC nuclei. Consistent with this, RESF1 has been reported to localise to nuclei when overexpressed in ESCs (Fukuda et al., 2018), a finding we confirm here using ESCs carrying epitope tagged endogenous *Resf1* loci.

A previous study showed that *Resf1^-/-^* ESCs could be maintained in serum/LIF culture (Fukuda et al., 2018), suggesting that ESC self-renewal continues after *Resf1* deletion. We confirm this here using clonal ESC self-renewal assays performed at saturating LIF concentrations. However, *Resf1^-/-^* ESCs cultured at low LIF concentrations show reduced self-renewal compared to wild type ESCs. We have extended these observations by showing that RESF1 overexpression increased colony forming capacity of ESCs cultured in serum/LIF. Interestingly, the levels of the key pluripotency mRNAs *Nanog, Esrrb* and *Pou5f1* were reduced in *Resf1^-/-^* ESCs cultured in serum/LIF, an effect that was reversed by culture of *Resf1^-/-^* ESCs in LIF media supplemented with MEK and GSK3β inhibitors (2i). This suggests that RESF1 causes ESCs to become either less sensitive to MEK or more responsive to the positive effects of Wnt or LIF signalling. This is consistent with the results from the colony forming assays. Together, these results suggest that RESF1 has a positive effect on ESC self-renewal.

As the only previously reported function of RESF1 is to silence endogenous retroviruses (ERVs), the different effects of *Resf1* deletion in ESCs cultured in 2i/LIF and serum/LIF could relate to the different activities of ERVs in these conditions. ESCs express higher levels of IAPEy and ERVK mRNAs when cultured in 2i/LIF rather than serum/LIF (Hackett et al., 2017). Interestingly, ESCs express higher levels of *Resf1* mRNA when cultured in serum/LIF rather than 2i/LIF, suggestive of a reciprocal relationship. Therefore, RESF1 function may be less critical during 2i/LIF culture. However, further investigation is needed to determine basis of higher tolerance of ESCs cultured in 2i/LIF to RESF1 depletion and ERV activity.

RESF1 also interacts with the histone methyltransferase SETDB1. In *Resf1^-/-^* ESCs, a decrease in SETDB1 binding and trimethylated histone 3, lysine 9 (H3K9me3) at an integrated MSCV proviral reporter locus suggests that RESF1 may act by supporting chromatin binding of SETDB1 (Fukuda et al., 2018). Similarly to *Resf1* deletion, deletion of SETDB1 or its associated protein TRIM28 leads to downregulation of the key pluripotency genes *Pou5f1, Sox2* and *Nanog*, upregulation of differentiation markers and ESC differentiation (Bilodeau et al., 2009; Hu et al., 2009; Karimi et al., 2011; Rowe et al., 2010; Yuan et al., 2009). Interestingly, a recent report indicates that another SETDB1 partner YTHDC1 also silences retrotransposons in ESCs and is critically required for ESC self-renewal (Liu et al., 2021). As the functions of RESF1 and YTHDC1 overlap, this suggests that functions of YTHDC1 not shared by RESF1 are required for ESC self-renewal. Nevertheless, an interaction with SETDB1 may be required for RESF1 to support ESC identity.

A further indication that RESF1 may participate in the pluripotency GRN comes from the binding of RESF1 to NANOG and OCT4 (Gagliardi et al., 2013; van den Berg et al., 2010). To better understand the molecular function of RESF1 in ESC self-renewal, we examined the relationship between RESF1 and NANOG. We validated the interaction of RESF1 and NANOG in ESC nuclei. However, this interaction is not essential for NANOG to sustain LIF independent self-renewal as *Resf1^-/-^* ESCs self-renew in the absence of LIF following NANOG overexpression. Also, *Resf1^-/-^* ESCs overexpressing the NANOG downstream target ESRRB also sustained LIF-independent self-renewal. Therefore, RESF1 is not required for NANOG or ESSRB to sustain LIF-independent ESC self-renewal.

NANOG and RESF1 both function in the germline (Chambers et al., 2007; Dickinson et al., 2016; M. Zhang et al., 2018). Nanog is required to provide wild type numbers of PGCs and is able to confer germline specification in the absence of instructive external signals (Murakami et al., 2016), while *Resf1* deletion causes infertility in both male and female mice (Dickinson et al., 2016). RESF1 may function during late germline development (Hudson et al., 2005) as deletion of the RESF1 partner protein SETDB1 prevents ERV silencing in E13.5 PGCs and blocks germline development (Liu et al., 2014). However, *Resf1* mRNA was also detected in PGCs at E6.75 and at higher levels than in epiblast cells at this stage, suggestive of a possible role in PGC specification. Indeed, *Resf1^-/-^* cells differentiate into PGCLCs with a lower efficiency *in vitro*, as judged by SSEA1 and CD61 expression and express lower levels of the key PGC transcription factor *Blimp1*. This suggests that RESF1 functions early during PGC specification.

Our results suggests that RESF1 has a modest effect on ESC self-renewal, and plays a role in early germline specification. It will be interesting to investigate the contribution of RESF1 to germline development *in vivo* and to determine how RESF1 safeguards fertility. Our ESCs carrying endogenously labelled *Resf1* alleles may be a valuable molecular tool for deciphering RESF1 function.

## Materials and Methods

### Cell culture

All cell lines were derived from E14Tg2a ESCs (Hooper et al., 1987) and were routinely cultured on 0.1% gelatin coated plates in serum/LIF medium [Glasgow Minimum Essential Medium (Sigma-Aldrich; G5154), 10 % foetal calf serum, 1x L-glutamine (Invitrogen, 25030-024), 1x pyruvate solution (Invitrogen, 11360-039), 1x MEM nonessential amino acids (Invitrogen, 11140-036), 0.1 mM 2-mercaptoethanol (Gibco, 31350010) and 100 U/ml LIF (homemade)] at 37 °C, 7 % CO_2_. Cells were passaged every other day using trypsin solution [0.372 mg/ml EDTA (Sigma cat. E5134), 1% chicken serum (Sigma, cat C5405), 0.025% w/v trypsin (Invitrogen, cat 15090-046)].

For 2i/LIF culture, ESCs cultured in serum/LIF medium were adapted to serum-free N2B27 medium supplemented with 3 µM CHIR99021 (Cambridge Bioscience, cat. 1677-5), 0.4 µM PD0325901 (STEMCELL Technologies, cat. 72182) and 100 U/ml homemade LIF as described by (Hayashi and Saitou, 2013) for at least 3 passages. The N2B27 medium was prepared as described by (Hayashi and Saitou, 2013). Cells were passaged on plates pre-treated with 0.01% w/v poly-L-ornithine and coated with 10 ng/ml laminin (BD Biosciences, cat. 354232).

ESCs were differentiated into EpiSCs as previously described (Guo 2009). 3×10^4^ ESCs cultured in serum/LIF medium were plated on a gelatin-coated 6-well plate and cultured for one day. Cells were washed twice with PBS and further cultured in EpiSC medium made by supplementing N2B27 medium with 20 ng/ml bFGF (R&D Systems, cat. 233-FB-025/CF) and 20 ng/ml Activin A (PeproTech, cat. 120-14E). After 24 hours, cells were washed twice using PBS and dissociated by 200 µl TrypLE Express (Gibco, cat 12604013) for 2 min at 37 °C. Cells were further cultured on plates coated with 7.5 μg/ml Fibronectin (Sigma, cat. F1141) in EpiSCs medium. EpiSCs were analysed after 6 passages.

ESCs were differentiated into PGCLCs as previously described (Hayashi and Saitou, 2013; J. Zhang et al., 2018). ESCs cultured in 2i/LIF medium were treated with TrypLE Express (Gibco, cat 12604013) to obtain a single cell suspension. 1×10^5^ ESCs were plated on a 3.8 cm^2^ plate coated with human plasma fibronectin (Millipore, cat. no. FC010) and cultured in N2B27 medium supplemented with 12 ng/ml bFGF (R&D Systems, cat. 233-FB-025/CF), 20 ng/ml Activin A (PeproTech, cat. 120-14E) and 1% KSR (Gibco, cat. 10828028) for 44 hours. Cells were treated with TrypLE Express and resuspended in GK15 medium [GMEM (Sigma, cat. G5154), 15% KSR (Invitrogen, cat. 10828-028), 1x nonessential amino acids (Invitrogen cat. 11140-036), 1 mM sodium pyruvate (Invitrogen, cat. 11360-039), 2 mM L-glutamine (Invitrogen, cat. 25030-024), 1:100 penicillin-streptomycin (Invitrogen, cat. 15070) and 0.1 mM 2-mercaptoethanol (Invitrogen, cat. No. 21985-023)] with freshly added 50 ng/ml Bmp4 (R&D Systems, cat. 314-BP-010), 50 ng/ml Bmp8a (R&D Systems, cat. 1073-BP-010), 2 ng/ml SCF (R&D Systems, cat. 455-MC-010), 500 ng/ml EGF (R&D Systems, cat. 2028-EG-010) and 1000 U/ml ESGRO (Millipore, ESG1106) to obtain a single cell suspension (1.5×10^5^ cells/ml). Cell suspension (100 µl/well) was added to 96 U-bottom well plates (Greiner-Bio, cat. 650970) and incubated at 37 °C, 5% CO_2_ for 4 days.

EpiSCs were prepared by ESC differentiation as previously described (Guo 2009). 3×10^4^ ESCs cultured serum/LIF medium were plated on a 6-well plate coated with 0.1 % gelatin and cultured for one day. Cells were washed twice with PBS and further cultured in N2B27 medium supplemented with 20 ng/ml bFGF (R&D Systems, cat. 233-FB-025/CF) and 20 ng/ml Activin A (PeproTech, cat. 120-14E). After 24 hours, cells were washed twice with PBS and dissociated using 200 µl TrypLE Express (Gibco, cat 12604013) for 2 min at 37 °C. Cells were further cultured on plates coated with 7.5 μg/ml Fibronectin (Sigma, cat. F1141) in the same medium. EpiSCs were analysed after 6 passages.

### Colony forming assays

Clonal assays were performed as described previously (Chambers et al., 2003). Briefly, 600 cells in a single cell suspension were plated per 9.5 cm^2^ and cultured for 6 days in serum medium and indicated concentrations of LIF (homemade). Formed colonies were fixed and stained for alkaline phosphatase using Leukocyte Alkaline Phosphatase Kit (Sigma; 86R-1KT) according to manufacturer’s instructions.

### Molecular cloning of *Resf1*

To clone the *Resf1* coding sequence, total RNA extract was prepared from E14Tg2a ESCs using RNeasy Mini Kit (Qiagen, 74104). The first cDNA strand was synthesized using oligo d(T) primers and Superscript III (Invitrogen, 18080093). *Resf1* open reading frame was PCR amplified from the prepared cDNA in two overlapping parts. The primer overhangs introduced sites complementary to pBlueScript plasmid to the 5’ and 3’ ends of the coding sequence. In addition, a triple flag tag and a glycine linker were inserted in front of the *Resf1* start codon. Both PCR products were subcloned into Blunt-TOPO vector using Zero Blunt TOPO PCR Cloning Kit (Invitrogen, cat. K2800-20SC) and verified by Sanger sequencing. Two parts of the *Resf1* open reading frame were PCR amplified from the TOPO vectors and cloned into XhoI, NotI digested pPyPPGK plasmid (Chambers et al., 2003) using homemade Gibson assembly mix (50 mM Tris-HCl pH 8.0, 5 mM MgCl_2_, 0.1 mM dNTPs, 25 mU/µl Phusion polymerase and 8 mU/µl T5 exonuclease).

### Transient episomal transfection

E14/T ESCs (Chambers et al., 2003) were transfected with pPyCAG-Flag_3_-Resf1-PGK-PURO, pPyCAG-Flag3-Nanog-PGK-PURO or pPyCAGPP (Chambers et al., 2003) plasmids using Lipofectamine 3000 (Thermo Fisher, cat. no. L3000001). Transfected cells were cultured overnight in serum/LIF medium at 37 °C, 7% CO_2_. Cells were washed with PBS, dissociated using trypsin solution (as above) and analysed by a clonal assay (as above) in the presence of puromycin.

### Stable integration of transgenes into ESCs

To assess self-renewal of ESCs with ectopic expression of NANOG, ESRRB or DsRed transgenes, linearised and purified pPyCAG-(Flag)_3_-Nanog-IRES-Puro, pPyCAG-(Flag)_3_-Esrrb-IRES-Puro and pPyCAG-DsRed-IRES-Puro plasmids (Chambers et al., 2003; Festuccia et al., 2012) were used to transfect *Resf1^-/-^* and E14Tg2a ESCs using Lipofectamine 3000 (Invitrogen cat. L3000001). Transfected cells were passaged every other day in serum/LIF medium supplemented with puromycin for 6 passages. Selected populations of cells were analysed by a colony forming assay in the presence of puromycin as described above.

### Deletion of *Resf1* gene in ESCs

To delete *Resf1*, two sets of four gRNAs (Supplementary table 1) targeting each end of the *Resf1* locus were cloned into eSpCas9(1.1)-T2A-eGFP and eSpCas9(1.1)-T2A-mCherry (Adgene #71814) plasmids, respectively, as previously described (Ran et al., 2013). E14Tg2a ESCs were transfected with the eSpCas9 plasmids using lipofectamine 3000 (Invitrogen cat. L3000001). After 24 hours, single cells expressing GFP and mCherry were isolated using fluorescence-activated cell sorting and expanded in serum/LIF medium. The isolated clonal cell lines were genotyped using primer pair A amplifying intergenic *Resf1* region and primer pair B which flanks the *Resf1* gene (Supplementary table 1). To verify *Resf1* deletion in individual alleles, PCR products were subcloned into Blunt-TOPO vector using Zero Blunt TOPO PCR Cloning Kit (Invitrogen, cat. K2800-20SC) and sequenced using Sanger sequencing.

### Endogenous tagging of *Resf1*

To create endogenously tagged *Resf1* cell lines, a donor vector was constructed by cloning 1kb 5’ and 3’homology arms together with the (v5)_3_-Stop-IRES-eGFP-STOP-loxP insert cassette into pBlueScript plasmid. The homology arms were amplified from E14Tg2a genomic DNA by Q5 polymerase (NEB, cat. no. M0491). The primers used introduced overhangs complementary to the insert cassette on the one side and pBlueScript on the other side. Homology arms and the insert cassette were cloned into EcoRI-HF (NEB cat. no. R3101S) cut pBlueScript using home-made Gibson assembly mix (described above). E14Tg2a ESCs were transfected with 1 µg of the donor vector as well as 1µg of the eSpCas9(1.1)-T2A-Puro plasmid coding for a single gRNA targeting the *Resf1* stop codon (Supplementary table 1) using Lipofectamine 3000 (Invitrogen cat. L3000001). The next day, culture medium was replaced with serum/LIF medium supplemented with puromycin and cultured for one more day. Cells were washed twice with PBS and dissociated using trypsin solution (see above). Single cells with high GFP signal were isolated and expanded. DNA from the individual clonal cell lines was isolated using DNeasy Blood & Tissue Kits (QIAGEN, cat. 69504) and genotyped using Rv5 genotyping primer pair (Supplementary table 1).

### RT-qPCR

ESCs were washed with PBS and lysed in 350 μl RLT buffer (QIAGEN) supplemented with 2-mercaptoethanol (1.4 M; Sigma, cat. no. M6250). RNA was extracted using RNeasy Plus Mini kit (QIAGEN, cat. 74136) according to the instructions. RNA was resuspended in nuclease-free H_2_O and stored at -80 °C. Purified RNA was reverse transcribed using SuperScript III (Invitrogen, cat. 18080085) according to the recommended protocol. Briefly, 0.2 - 1 μg of RNA was mixed with 50 ng of random hexamers (Invitrogen, cat. N8080127) and 1 μl of 10 mM dNTPs (Invitrogen, cat. 10297018) in the final volume of 11 μ The mix was incubated at 65 °C for 5 min followed by 2 min in ice. Next, 2 μl of 100 mM DTT, 20 U of RNAseOUT (Invitrogen, cat. 10777019) and 100 U of SuperScript III reverse transcriptase (Invitrogen, cat. 18080044) were added. The reaction mix was adjusted to the final volume of 20 μl with nuclease-free H_2_O and incubated 10 min at 25 °C, 1 h at 50 °C and 15 min at 70 °C. cDNA was diluted 1:10 (v/v) before quantification. The qPCR reaction mix was prepared by mixing 5 μl of the cDNA, 4.5 μl of Takyon™ SYBR 2X qPCR Mastermix (Eurogentec) and 0.5 μl of a primer pair mix (10 mM each) (Supplementary table 1) in a 384-well plate in duplicates. Specificity of the used primers was determined from a melting curve and their efficiency (>90%) by a linear regression.

### Co-immunoprecipitation

The E14/T ESCs (6×10^6^) were transfected with pPyCAG-(HA)_3_-Nanog-IRES-Puro and pPyCAG-(Flag)_3_-Resf1-IRES-Puro using Lipofectamine 3000 (Invitrogen cat. L3000001). In parallel, E14/T ESCs were transfected with only pPyCAG-(HA)_3_-Nanog-IRES-Puro plasmid as a negative control. The day after the transfection, medium was replaced with fresh serum/LIF medium supplemented with G418 and puromycin and cultured for one day. Cells were detached by trypsin, washed with PBS and collected. Cells were burst in a hypotonic buffer (5 mM Pipes pH 8, 85 mM KCl) with freshly added 0.5% NP-40 and 1x Protease Inhibitor Cocktail (Roche, cat. 4693159001) for 20 min in ice. Nuclei were collected by centrifugation (1,500 rpm, 5 min, 4 C), resuspended in 1 ml of the NE buffer (20 mM HEPES pH 7.6, 350 mM KCl, 0.2 mM EDTA pH 8, 1.5 mM MgCl_2_, 20 % Glycerol) with freshly added 0.2% NP-40, 0.5 mM DTT and 1x Protease Inhibitor Cocktail (Roche, cat. no. 4693159001) and transferred into NoStick microtubes (Alpha Laboratories, Cat. LW2410AS). Nuclei were lysed for 1 h at 4 °C in a presence of 150 U of Benzonase nuclease (Novagen, cat. no. 71206) while rotating. Nuclear lysates were cleared by centrifugation (13,300 rpm, 30 min, 4 C) and transferred into clean NoStick tubes. Input control fractions (5%) were separated. Anti-FLAG M2 affinity beads slurry (30 µl, Sigma cat. no. A2220) was washed three times with PBS and resuspended in the original volume of the NE buffer. Thirty microlitres of the beads were added to the remaining nuclear extracts and incubated for 2 h at room temperature on a wheel. The beads were washed three times with PBS using magnet. Proteins were eluted by boiling the beads for 5 min in 30 μl of 1x LDS Sample Buffer (Invitroge, cat. no. B0008) supplemented with 250 mM DTT. The elution was repeated, and the two fractions were combined. The input and immunoprecipitated samples were analysed by SDS-PAGE and immunoblot.

### SDS-PAGE and immunoblotting

The protein extracts were mixed with 4X Bolt™ LDS Sample Buffer (Invitrogen, cat. no. B0008) and 250 mM DTT and boiled for 5 min. Samples together with SeeBlue™ Plus2 pre-stained protein standard (Invitrogen, cat. no. LC5925) were loaded on a Bolt™ 12% Bis-Tris Plus Gel (Life Biosciences, cat. no. NW00122BOX) and run at constant 180 V in a Bolt™ MES SDS running buffer (Invitrogen, cat. no. B0002). The proteins were transferred onto a nitrocellulose membrane in transfer buffer (25 mM Tris-HCl pH 8, 0.21 M Glycine, 10% methanol) at constant 180 mA over night at 4 °C or at 380 mA for 70 min in ice. The membrane was blocked in 10% skimmed milk resuspended in PBS and supplemented with 0.1% Tween-20 (PBS-T) for 1 h or overnight. Primary antibodies (Supplementary table 1) were diluted in 5 ml of 5% milk in PBS-T and added on the membrane. The membrane was stained for at least 1 hour at the room temperature while swirling. The membrane was washed four times for 5 min with PBS-T. The secondary antibodies (Supplementary table 1) were diluted in 5 ml of 5% milk in PBS-T and incubated with the membrane for at least 1 h at the room temperature. The membrane was washed four times for 5 min in PBS-T. The membrane was incubated with Pierce™ ECL Western Blotting Substrate (Invitrogen, cat. no. 32106) for 2 min at room temperature if an antibody with conjugated horse radish peroxidase was used.

### Immunostaining

Cells were washed with PBS and fixed with 4% paraformaldehyde at room temperature for 10 min. Cells were washed with PBS and permeabilised in 0.3% v/v Triton X-100 in PBS for 15 min at room temperature. The solution was discarded, and cells were blocked in 0.1% Triton X-100, 3% v/v donkey serum in PBS for 1 h at room temperature. Fixed and permeabilised cells were incubated with primary antibodies (Supplementary table 1) diluted in blocking buffer over night at 4 °C. Cells were washed four times with PBS containing 0.1% Triton X-100 and incubated with the fluorophore-labelled secondary antibodies diluted (1:1,000 v/v) in blocking buffer for 1 h at room temperature. DNA was stained with 4’,6-diamidino-2-phenylindole (DAPI; 1:2,000 dilution in PBS) for 5-10 min at room temperature. DAPI was washed once with PBS for 5 min and samples were stored in a mounting solution at 4 °C in dark. Samples were analysed using SP8 Lightning confocal microscope (Leica).

### Flow-cytometry

To quantify SSEA1 and CD61 expression, cell aggregates were collected washed with PBS and dissociated in 0.1% Trypsin solution at 37 °C for 10 min. Trypsin was neutralised with serum medium and passed through a cell strainer. A small proportion of each analysed suspension was combined in a separate tube for control samples. Cells were centrifuged (1,100 rpm, 3 min) and resuspended in 100 μl of serum medium supplemented with 0.5 μl SSEA-I (BioLegend, cat. no. 125607) and 0.15 μl PE-CD61 antibodies (BioLegend, cat. no. 104307). The negative control was resuspended in 300 μl of the serum medium and divided 100 μl fractions. SSEA-I and CD61 antibodies were added to one fraction resulting in two single-stain control samples and one negative control. Cells were incubated for 15 min at room temperature in dark and washed twice with PBS. Cell were resuspended in 200 μl of 2% KSR in PBS and analysed on a 5 laser LSR Fortessa analyser (BD Biosciences). Single cells were gated based on forward and side scatters. Live cells were gated based on DAPI signal and the SSEA-I, CD61 double positive population was gated based on the negative and single stain controls.

### Analysis of single cell RNA sequencing data

Single cell RNA sequencing data of mouse embryo were obtained using R package MouseGastrulationData (Griffiths and Lun, 2020). The number of individual cells expressing *Resf1* (log normalised counts > 0) was determined in epiblast and PGCs at indicated embryonic stages. *Resf1* expression in epiblast and PGCs at different embryonic stages was visualised by plotting log normalised counts of *Resf1* in cells with *Resf1* log normalised counts > 0.

## Acknowledgements

We thank members of the Chambers lab for discussions and comments on the manuscript. This work was funded by a Medical Research Council (UK) grant to I.C. (MR/T003162/1), M.V. was supported by a Principal’s Career Development Scholarship from the University of Edinburgh.

---

**Figure S1:**
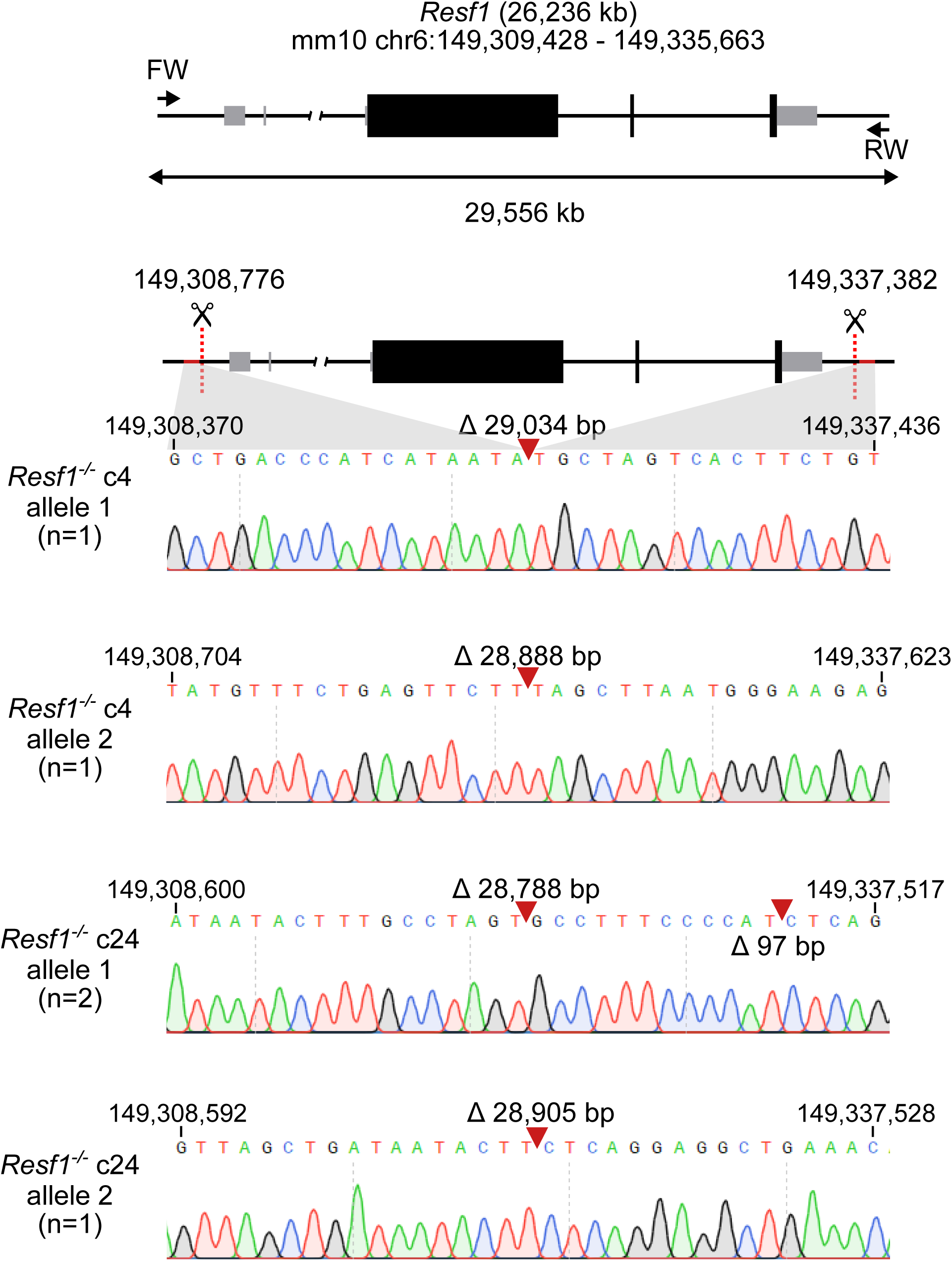
Sequence analysis of putative *Resf1^-/-^* ES cell lines. Cartoon of the *Resf1* locus. *Resf1* size and genome coordinates (mm10 genome assembly) are shown at the top. Thick black boxes represent coding exons, thin grey boxes represent non-coding exons, lines represent introns and non-transcribed regions. Forward (FW) and reverse (RV) primers used to analyse deletions of *Resf1* are indicated as arrows. The distance between FW and RV primers is shown by the double arrow. *S*ites 5’ of *Resf1* start codon and 3’ from *Resf1* polyadenylation signal targeted by CRISPR/Cas9 are shown as red dotted lines. Deletion of *Resf1* was validated in putative *Resf1^-/-^* clones 4 (c4) and 24 (c24) by PCR using FW and RV primers. Sequence tracks of the amplified region in individual alleles of *Resf1^-/-^* c4 and c24 cell lines. Complementary coordinates of the first and the last shown residue are shown. Red arrows indicate deletion events with indicated size of the deletion.

**Figure S2:**
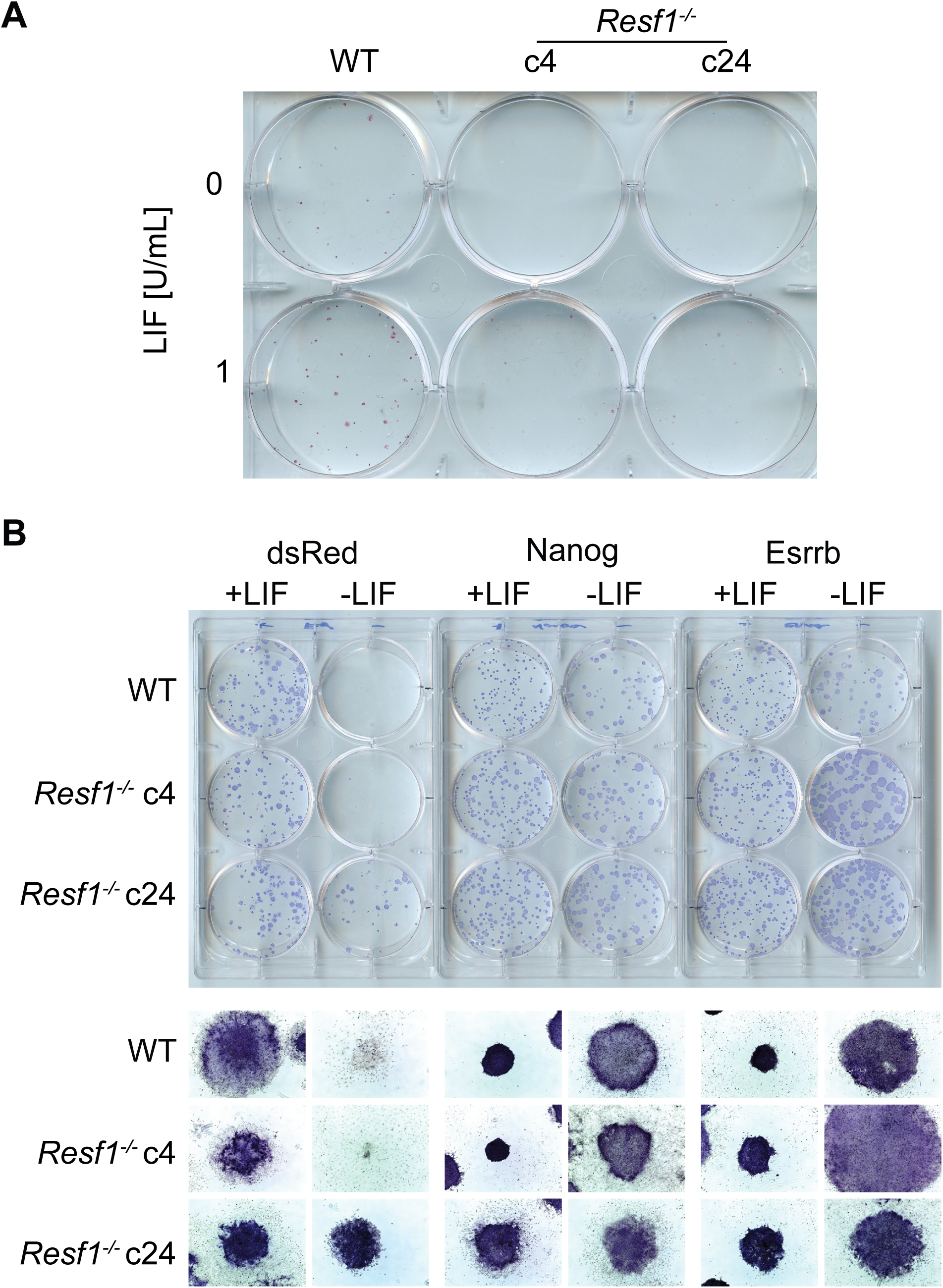
Self-renewal of *Resf1^-/-^* ESCs at low LIF concentrations and capacity of NANOG and ESRRB to confer LIF-independent self-renewal in *Resf1^-/-^* ESCs. **A)** Photography of colonies formed by wild type (WT) and *Resf1^-/-^* clonal cell lines (c4, c24) after 6-day culture in serum medium supplemented with no (0) or 1 U/ml LIF. Colonies were stained for alkaline phosphatase. **B)** Images of colonies stained for alkaline phosphatase of WT or *Resf1^-/-^* ESCs stably expressing dsRed, Flag-Nanog or Flag-Esrrb transgenes cultured in the presence (+) or absence (-) of LIF (100 U/ml) for 8 days.

**Figure S3:**
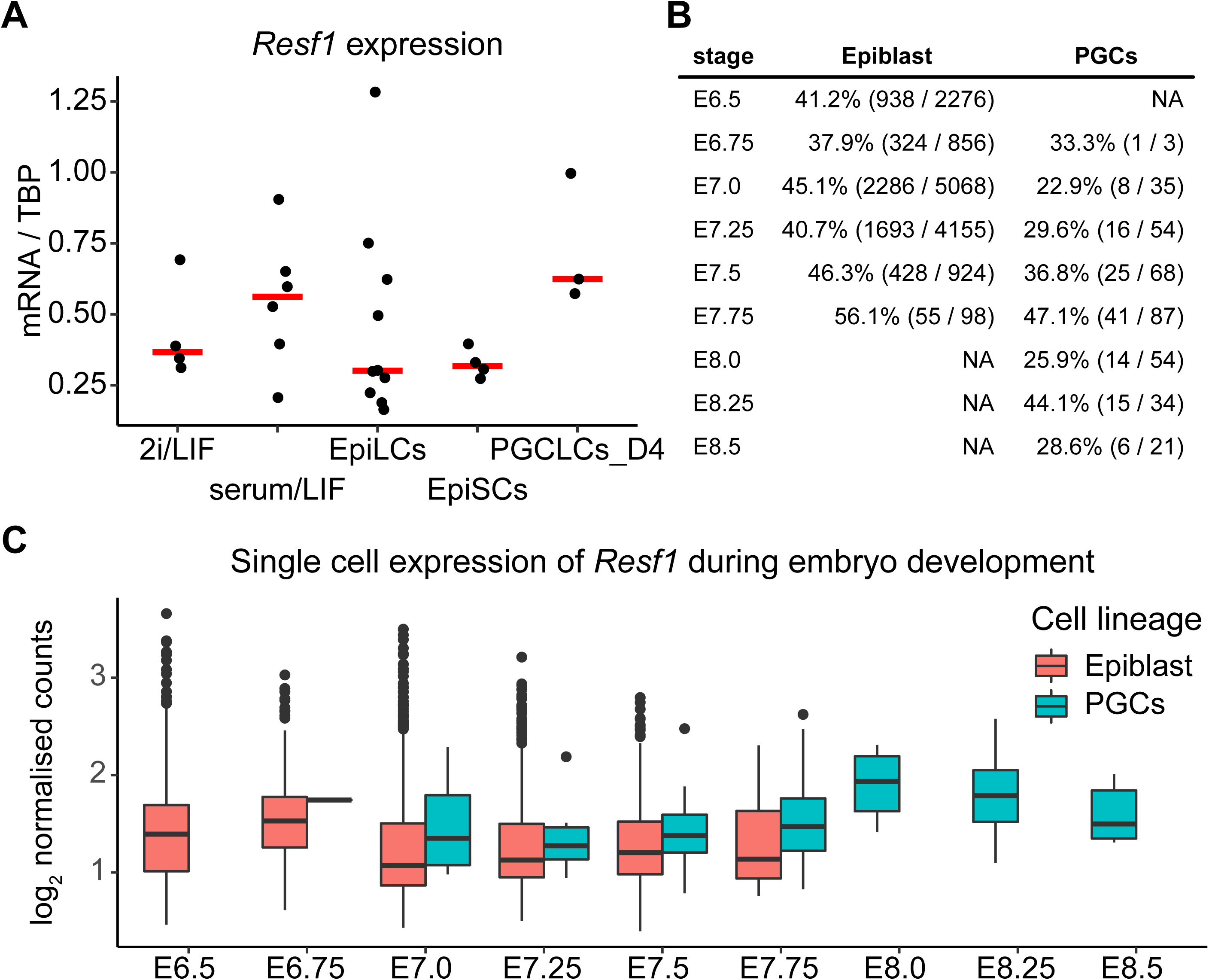
Resf1 expression in epiblast and primordial germ cells. **A)** RT-qPCR analysis of *Resf1* mRNA levels normalised to TBP mRNA in E14Tg2a embryonic stem cells cultured in 2i/LIF or serum/LIF medium, epiblast-like cells (EpiLCs), epiblast stem cells (EpiSCs) and day 4 primordial germ cell-like cells (PGCLCs_D4). Individual data points are shown as well as the median expression levels (red bar). **B)** and **C)** *Resf1* expression in single cell RNA sequencing dataset of mouse embryo at the indicated embryonic (E) stages. **B)** Summary of *Resf1* mRNA expression in epiblast and primordial germ cells (PGCs) between embryonic stages E6.5 and E8.5. The proportion of single cells expressing *Resf1* at individual stages is shown. **C)** Boxplots showing *Resf1* mRNA expression levels in PGCs and epiblast cells from B. Box represents interquartile range; horizontal lines are medians and points are outliers.

**Figure S4:**
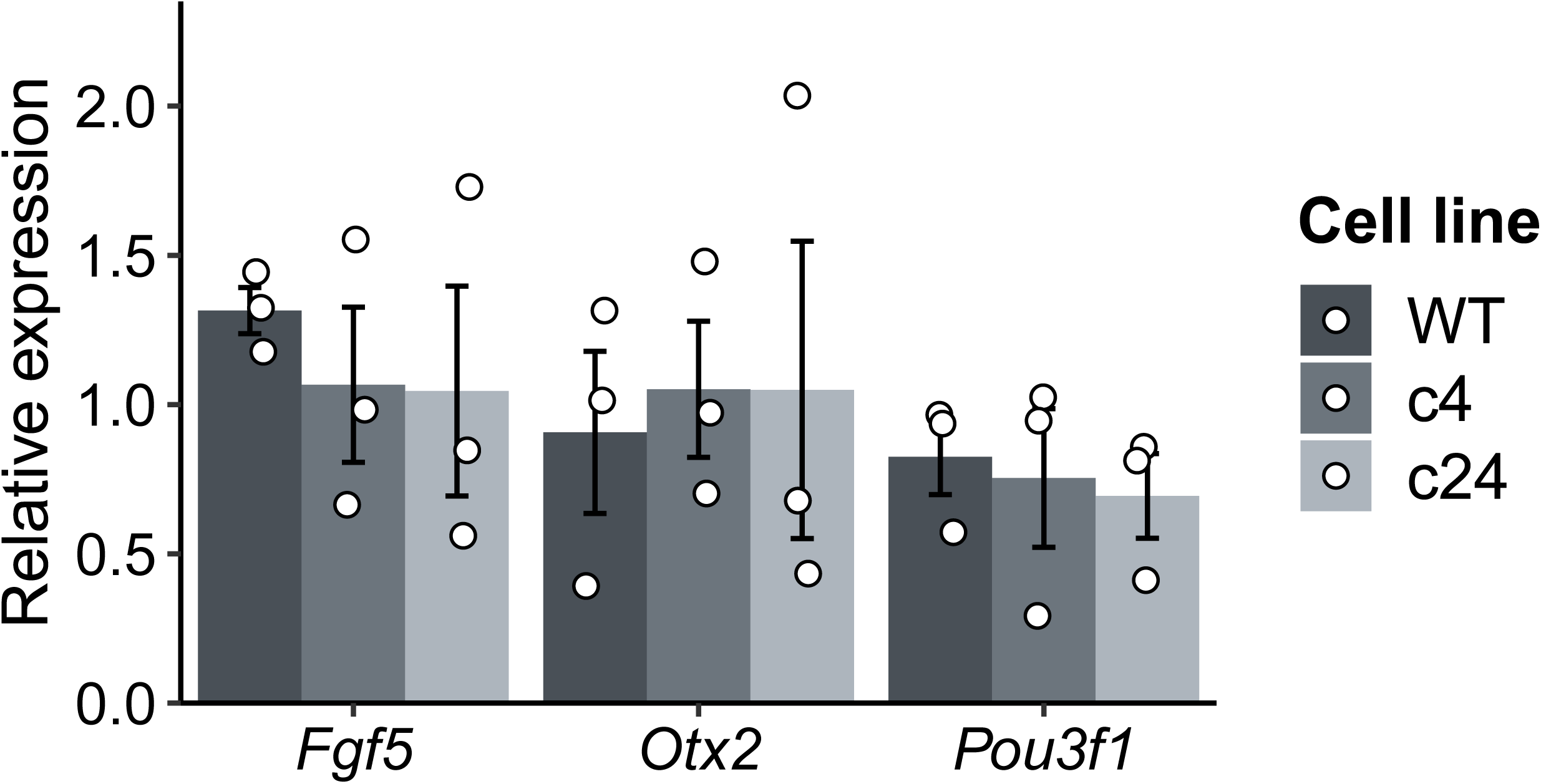
Expression of EpiLC markers in *Resf1^-/-^* EpiLCs. mRNA levels of the indicated transcripts were measured in wild type (WT) and *Resf1*^-/-^ EpiLCs (c4 and c24) by RT-qPCR. Bars and whiskers represent mean ± SD (n = 3). Scatter plots represent individual data points.

**Figure S5:**
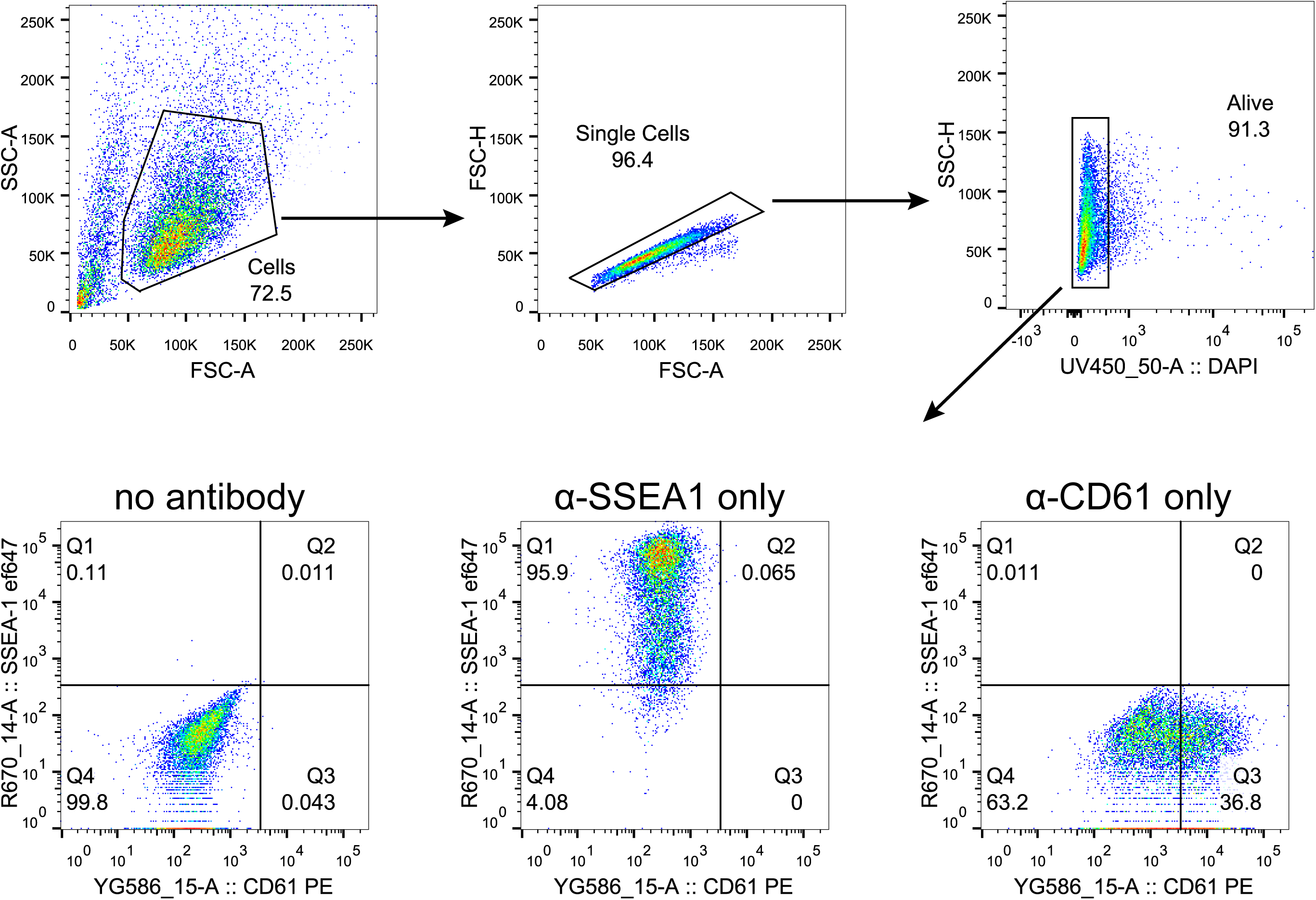
Gating strategy for quantification of PGCLC population. Representative scatter plots indicating gating strategy to quantify PGCLC population marked by co-expression of SSEA1 and CD61. Gating for live cells was done using DAPI. Samples without any antibody staining and samples stained with a single antibody were used to gate SSEA1+CD61+ population as shown.

## References

1. Bilodeau, S., Kagey, M.H., Frampton, G.M., Rahl, P.B., Young, R.A., 2009. SetDB1 contributes to repression of genes encoding developmental regulators and maintenance of ES cell state. Genes Dev. 23, 2484–2489. https://doi.org/10.1101/gad.1837309

2. Chambers, I., Colby, D., Robertson, M., Nichols, J., Lee, S., Tweedie, S., Smith, A., 2003. Functional Expression Cloning of Nanog, a Pluripotency Sustaining Factor in Embryonic Stem Cells. Cell 113, 643–655. https://doi.org/10.1016/S0092-8674(03)00392-1

3. Chambers, I., Silva, J., Colby, D., Nichols, J., Nijmeijer, B., Robertson, M., Vrana, J., Jones, K., Grotewold, L., Smith, A., 2007. Nanog safeguards pluripotency and mediates germline development. Nature 450, 1230–1234. https://doi.org/10.1038/nature06403

4. Dickinson, M.E., Flenniken, A.M., Ji, X., Teboul, L., Wong, M.D., White, J.K., Meehan, T.F., Weninger, W.J., Westerberg, H., Adissu, H., Baker, C.N., Bower, L., Brown, J.M., Caddle, L.B., Chiani, F., Clary, D., Cleak, J., Daly, M.J., Denegre, J.M., Doe, B., Dolan, M.E., Edie, S.M., Fuchs, H., Gailus-Durner, V., Galli, A., Gambadoro, A., Gallegos, J., Guo, S., Horner, N.R., Hsu, C.-W., Johnson, S.J., Kalaga, S., Keith, L.C., Lanoue, L., Lawson, T.N., Lek, M., Mark, M., Marschall, S., Mason, J., McElwee, M.L., Newbigging, S., Nutter, L.M.J., Peterson, K.A., Ramirez-Solis, R., Rowland, D.J., Ryder, E., Samocha, K.E., Seavitt, J.R., Selloum, M., Szoke-Kovacs, Z., Tamura, M., Trainor, A.G., Tudose, I., Wakana, S., Warren, J., Wendling, O., West, D.B., Wong, L., Yoshiki, A., Wurst, W., MacArthur, D.G., Tocchini-Valentini, G.P., Gao, X., Flicek, P., Bradley, A., Skarnes, W.C., Justice, M.J., Parkinson, H.E., Moore, M., Wells, S., Braun, R.E., Svenson, K.L., Angelis, M.H. de, Herault, Y., Mohun, T., Mallon, A.-M., Henkelman, R.M., Brown, S.D.M., Adams, D.J., Lloyd, K.C.K., McKerlie, C., Beaudet, A.L., Bućan, M., Murray, S.A., 2016. High-throughput discovery of novel developmental phenotypes. Nature 537, 508–514. https://doi.org/10.1038/nature19356

5. Festuccia, N., Osorno, R., Halbritter, F., Karwacki-Neisius, V., Navarro, P., Colby, D., Wong, F., Yates, A., Tomlinson, S.R., Chambers, I., 2012. Esrrb Is a Direct Nanog Target Gene that Can Substitute for Nanog Function in Pluripotent Cells. Cell Stem Cell 11, 477–490. https://doi.org/10.1016/j.stem.2012.08.002

6. Fukuda, K., Okuda, A., Yusa, K., Shinkai, Y., 2018. A CRISPR knockout screen identifies SETDB1-target retroelement silencing factors in embryonic stem cells. Genome Res. https://doi.org/10.1101/gr.227280.117

7. Gagliardi, A., Mullin, N.P., Ying Tan, Z., Colby, D., Kousa, A.I., Halbritter, F., Weiss, J.T., Felker, A., Bezstarosti, K., Favaro, R., Demmers, J., Nicolis, S.K., Tomlinson, S.R., Poot, R.A., Chambers, I., 2013. A direct physical interaction between Nanog and Sox2 regulates embryonic stem cell self-renewal. The EMBO Journal 32, 2231–2247. https://doi.org/10.1038/emboj.2013.161

8. Griffiths, J., Lun, A., 2020. MouseGastrulationData: Single-Cell Transcriptomics Data across Mouse Gastrulation and Early Organogenesis.

9. Hackett, J.A., Kobayashi, T., Dietmann, S., Surani, M.A., 2017. Activation of Lineage Regulators and Transposable Elements across a Pluripotent Spectrum. Stem Cell Reports 8, 1645–1658. https://doi.org/10.1016/j.stemcr.2017.05.014

10. Hayashi, K., Ohta, H., Kurimoto, K., Aramaki, S., Saitou, M., 2011. Reconstitution of the Mouse Germ Cell Specification Pathway in Culture by Pluripotent Stem Cells. Cell 146, 519–532. https://doi.org/10.1016/j.cell.2011.06.052

11. Hayashi, K., Saitou, M., 2013. Generation of eggs from mouse embryonic stem cells and induced pluripotent stem cells. Nature Protocols 8, 1513–1524. https://doi.org/10.1038/nprot.2013.090

12. Hooper, M., Hardy, K., Handyside, A., Hunter, S., Monk, M., 1987. HPRT-deficient (Lesch–Nyhan) mouse embryos derived from germline colonization by cultured cells. Nature 326, 292–295. https://doi.org/10.1038/326292a0

13. Hu, G., Kim, J., Xu, Q., Leng, Y., Orkin, S.H., Elledge, S.J., 2009. A genome-wide RNAi screen identifies a new transcriptional module required for self-renewal. Genes Dev. 23, 837–848. https://doi.org/10.1101/gad.1769609

14. Hudson, Q.J., Smith, C.A., Sinclair, A.H., 2005. Conserved expression of a novel gene during gonadal development. Developmental Dynamics 233, 1083–1090. https://doi.org/10.1002/dvdy.20397

15. Karimi, M.M., Goyal, P., Maksakova, I.A., Bilenky, M., Leung, D., Tang, J.X., Shinkai, Y., Mager, D.L., Jones, S., Hirst, M., Lorincz, M.C., 2011. DNA Methylation and SETDB1/H3K9me3 Regulate Predominantly Distinct Sets of Genes, Retroelements, and Chimeric Transcripts in mESCs. Cell Stem Cell 8, 676–687. https://doi.org/10.1016/j.stem.2011.04.004

16. Kinoshita, M., Smith, A., 2018. Pluripotency Deconstructed. Development, Growth & Differentiation 60, 44–52. https://doi.org/10.1111/dgd.12419

17. Liu, J., Gao, M., He, J., Wu, K., Lin, S., Jin, L., Chen, Y., Liu, H., Shi, J., Wang, X., Chang, L., Lin, Y., Zhao, Y.-L., Zhang, X., Zhang, M., Luo, G.-Z., Wu, G., Pei, D., Wang, J., Bao, X., Chen, J., 2021. The RNA m 6 A reader YTHDC1 silences retrotransposons and guards ES cell identity. Nature 591, 322–326. https://doi.org/10.1038/s41586-021-03313-9

18. Liu, S., Brind’Amour, J., Karimi, M.M., Shirane, K., Bogutz, A., Lefebvre, L., Sasaki, H., Shinkai, Y., Lorincz, M.C., 2014. Setdb1 is required for germline development and silencing of H3K9me3-marked endogenous retroviruses in primordial germ cells. Genes Dev. 28, 2041–2055. https://doi.org/10.1101/gad.244848.114

19. Martello, G., Smith, A., 2014. The Nature of Embryonic Stem Cells. Annual Review of Cell and Developmental Biology 30, 647–675. https://doi.org/10.1146/annurev-cellbio-100913-013116

20. Mitsui, K., Tokuzawa, Y., Itoh, H., Segawa, K., Murakami, M., Takahashi, K., Maruyama, M., Maeda, M., Yamanaka, S., 2003. The Homeoprotein Nanog Is Required for Maintenance of Pluripotency in Mouse Epiblast and ES Cells. Cell 113, 631–642. https://doi.org/10.1016/S0092-8674(03)00393-3

21. Murakami, K., Guenesdogan, U., Zylicz, J.J., Tang, W.W.C., Sengupta, R., Kobayashi, T., Kim, S., Butler, R., Dietmann, S., Surani, M.A., 2016. NANOG alone induces germ cells in primed epiblast in vitro by activation of enhancers. Nature 529, 403–+. https://doi.org/10.1038/nature16480

22. Nichols, J., Smith, A., 2009. Naive and Primed Pluripotent States. Cell Stem Cell 4, 487–492. https://doi.org/10.1016/j.stem.2009.05.015

23. Pantier, R., Tatar, T., Colby, D., Chambers, I., 2019. Endogenous epitope-tagging of Tet1, Tet2 and Tet3 identifies TET2 as a naïve pluripotency marker. Life Sci Alliance 2. https://doi.org/10.26508/lsa.201900516

24. Pijuan-Sala, B., Griffiths, J.A., Guibentif, C., Hiscock, T.W., Jawaid, W., Calero-Nieto, F.J., Mulas, C., Ibarra-Soria, X., Tyser, R.C.V., Ho, D.L.L., Reik, W., Srinivas, S., Simons, B.D., Nichols, J., Marioni, J.C., Göttgens, B., 2019. A single-cell molecular map of mouse gastrulation and early organogenesis. Nature 566, 490–495. https://doi.org/10.1038/s41586-019-0933-9

25. Ran, F.A., Hsu, P.D., Wright, J., Agarwala, V., Scott, D.A., Zhang, F., 2013. Genome engineering using the CRISPR-Cas9 system. Nature Protocols 8, 2281–2308. https://doi.org/10.1038/nprot.2013.143

26. Rowe, H.M., Jakobsson, J., Mesnard, D., Rougemont, J., Reynard, S., Aktas, T., Maillard, P.V., Layard-Liesching, H., Verp, S., Marquis, J., Spitz, F., Constam, D.B., Trono, D., 2010. KAP1 controls endogenous retroviruses in embryonic stem cells. Nature 463, 237–240. https://doi.org/10.1038/nature08674

27. Silva, J., Nichols, J., Theunissen, T.W., Guo, G., van Oosten, A.L., Barrandon, O., Wray, J., Yamanaka, S., Chambers, I., Smith, A., 2009. Nanog Is the Gateway to the Pluripotent Ground State. Cell 138, 722–737. https://doi.org/10.1016/j.cell.2009.07.039

28. Silva, J., Smith, A., 2008. Capturing Pluripotency. Cell 132, 532–536. https://doi.org/10.1016/j.cell.2008.02.006

29. van den Berg, D.L.C., Snoek, T., Mullin, N.P., Yates, A., Bezstarosti, K., Demmers, J., Chambers, I., Poot, R.A., 2010. An Oct4-centered protein interaction network in embryonic stem cells. Cell Stem Cell 6, 369–381. https://doi.org/10.1016/j.stem.2010.02.014

30. Ying, Q.-L., Wray, J., Nichols, J., Batlle-Morera, L., Doble, B., Woodgett, J., Cohen, P., Smith, A., 2008. The ground state of embryonic stem cell self-renewal. Nature 453, 519–523. https://doi.org/10.1038/nature06968

31. Yuan, P., Han, J., Guo, G., Orlov, Y.L., Huss, M., Loh, Y.-H., Yaw, L.-P., Robson, P., Lim, B., Ng, H.-H., 2009. Eset partners with Oct4 to restrict extraembryonic trophoblast lineage potential in embryonic stem cells. Genes Dev. 23, 2507–2520. https://doi.org/10.1101/gad.1831909

32. Zhang, J., Zhang, M., Acampora, D., Vojtek, M., Yuan, D., Simeone, A., Chambers, I., 2018. OTX2 restricts entry to the mouse germline. Nature 562, 595–599. https://doi.org/10.1038/s41586-018-0581-5

33. Zhang, M., Leitch, H.G., Tang, W.W.C., Festuccia, N., Hall-Ponsele, E., Nichols, J., Surani, M.A., Smith, A., Chambers, I., 2018. Esrrb Complementation Rescues Development of Nanog-Null Germ Cells. Cell Reports 22, 332–339. https://doi.org/10.1016/j.celrep.2017.12.060

